# Cell cycle length regulates heterochromatin reprogramming during early development in non-mammalian vertebrates

**DOI:** 10.1101/2024.01.23.576847

**Authors:** Hiroto S Fukushima, Takafumi Ikeda, Shinra Ikeda, Hiroyuki Takeda

**Affiliations:** Department of Biological Sciences, Graduate School of Science, The University of Tokyo, Tokyo 113-0033, Japan; Center for Integrative Medical Sciences, RIKEN, Yokohama 230-0045, Japan; Institute for Protein Dynamics, Kyoto Sangyo University, Kyoto 603-8555, Japan; Faculty of Life Sciences, Kyoto Sangyo University, Kyoto 603-8555, Japan

**Keywords:** Development, Epigenome, Heterochromatin, Mid-blastula transition, Reprogramming

## Abstract

Heterochromatin marks such as H3K9me3 undergoes global erasure and re-establishment after fertilization, and the proper reprogramming of H3K9me3 is essential for early development. Despite the widely conserved dynamics of heterochromatin reprogramming in invertebrates and non-mammalian vertebrates, previous studies have shown that the underlying mechanisms may differ between species. In this study, we investigated the molecular mechanism of H3K9me3 dynamics in medaka (Japanese killifish, *Oryzias latipes*) as a non-mammalian vertebrate model, and found that rapid cell cycle during the cleavage stages causes DNA replication-dependent passive erasure of H3K9me3. We also found that cell cycle slowing, toward the mid-blastula transition, permits increasing nuclear accumulation of H3K9me3 histone methyltransferase Setdb1, leading to the onset of H3K9me3 re-accumulation. We further demonstrated that cell cycle length in early development regulates H3K9me3 reprogramming in zebrafish and *Xenopus laevis* as well. Together with the previous studies in invertebrates, we propose that the cell cycle length-dependent mechanism for both global erasure and re-accumulation of H3K9me3 is widely conserved among rapid-cleavage species of non-mammalian vertebrates and invertebrates such as *Drosophila*, *C. elegans* and teleost fish.

## Introduction

Heterochromatin represses gene expression by physical folding and serves as epigenetic memory in cells, and the repressive histone modification, H3K9me3, plays a central role in the establishment and maintenance of heterochromatin (Allshire & Madhani, 2018; Padeken *et al*, 2022). After fertilization, cellular memories installed during gametogenesis undergo reprogramming, which allows embryonic cells to acquire pluripotency (Xia & Xie, 2020; Xu & Xie, 2018). Indeed, during this process, heterochromatin marks such as H3K9me3 are erased and re-installed (Mutlu *et al*, 2018, 2019; Seller *et al*, 2019; Laue *et al*, 2019; Fukushima *et al*, 2023; Bessler *et al*, 2010; Yuan & O’Farrell, 2016; Hontelez *et al*, 2015; Wang *et al*, 2018; Yu *et al*, 2022; Zhou *et al*, 2023). The proper reprogramming of H3K9me3 is essential for normal development. For example, previous studies of somatic cell nuclear transfer suggest that heterochromatin marks, if remain during early development, act as an epigenetic barrier to entry into normal development (Matoba *et al*, 2014; Chung *et al*, 2015; Jullien *et al*, 2017; Xu *et al*, 2023), and experimental removal of H3K9me3 leads to abnormal development (Fukushima *et al*, 2023). However, in spite of well-recognized dynamics and importance of heterochromatin reprogramming in early development, how this process is regulated remain elusive, in particular, in vertebrates.

In differentiated cells, H3K9me3 is deposited by histone methyltransferases Suv39h1/2 and Setdb1, and erased actively by enzymes (e.g. demethylase) or passively by DNA-replication-coupled dilution of histone modifications (Allshire & Madhani, 2018; Padeken *et al*, 2022). Although the timing and extent of heterochromatin reprogramming vary between species, they should be tightly regulated during early development because of their profound impacts on transcription and genome structure (Allshire & Madhani, 2018; Padeken *et al*, 2022). The vast majority of animals begin development with rapid cell divisions and exhibit the conserved dynamics of transcription and epigenetic reprogramming. Mammalian embryos, however, are characterized by a slow cell cycle, an early onset of transcription, and unique epigenetic reprogramming (Tadros & Lipshitz, 2009; Vastenhouw *et al*, 2019; Xia & Xie, 2020). In rapid-cleavage species such as *Drosophila*, *C. elegans*, and non-mammalian model vertebrates (Tadros & Lipshitz, 2009; Vastenhouw *et al*, 2019), H3K9 methylations are detected in germ cells, almost undetectable after fertilization, and detected again after several rounds of cleavage, indicating that H3K9me3 is subject to global erasure and re-accumulation (Mutlu *et al*, 2018, 2019; Seller *et al*, 2019; Laue *et al*, 2019; Fukushima *et al*, 2023; Bessler *et al*, 2010; Yuan & O’Farrell, 2016; Hontelez *et al*, 2015). In those species, the slowing of the cell cycle and zygotic genome activation (ZGA) coincide with the re-establishment of heterochromatin (Tadros & Lipshitz, 2009; Vastenhouw *et al*, 2019), raising the possibility that the two events are involved in the onset of H3K9me3 deposition in early development. However, despite the highly conserved dynamics of heterochromatin reprogramming among rapid-cleavage species, species-specific mechanisms have been proposed. In *Drosophila* and *C. elegans*, cell cycle slowing is a major mechanism that times the onset of H3K9me3 deposition by allowing nuclear accumulation of histone methyltransferase (Mutlu *et al*, 2018, 2019; Seller *et al*, 2019). On the other hand, Laue *et al*. demonstrated in zebrafish that ZGA-dependent clearance of the maternally provided chromatin remodeler, Smarca2, is a prerequisite for re-accumulation of H3K9me3 (Laue *et al*, 2019). Therefore, the underlying mechanism varies even between these rapid cleavage species. Furthermore, the mechanism that drives the erasure process is even less understood.

In this study, we address what drives the erasure of heterochromatin during cleavage and what determines the timing of heterochromatin re-establishment, mainly in medaka, Japanese killifish (*Oryzias latipes*). The medaka is another good non-mammalian model organism with a large evolutionary distance to other non-mammalian vertebrates (Furutani-Seiki & Wittbrodt, 2004; Takeda & Shimada, 2010), which has an extensive collection of epigenome data on reprogramming during early development (Nakamura *et al*, 2014, 2021; Fukushima *et al*, 2023). We demonstrate here that it is not ZGA but regulation of cell cycle length that mediates both the erasure and re-accumulation of H3K9me3 during early development in medaka. We also show that this holds true for zebrafish and *Xenopus* embryos. Taken together with the previous studies in *C. elegans* and *Drosophila* (Mutlu *et al*, 2018, 2019; Seller *et al*, 2019), our results suggest that cell cycle length regulation commonly and mainly causes global erasure and re-establishment of heterochromatin in rapid-cleavage species, and provide a new evolutionary perspective on the dynamics of epigenetic reprogramming in animal development.

## Results

### Passive erasure of heterochromatin during early cleavage stages in medaka

Previously, we revealed that H3K9me3 is observed in medaka embryos at the one-cell stage, but is almost undetectable at the 16-cell stage, except in telomeric regions (Fukushima *et al*, 2023). To examine how H3K9me3 disappears between the two stages, we performed immunofluorescence staining of H3K9me3 at several developmental points between the two stages (Fig 1A). We found that H3K9me3 was gradually decreased from the one-cell stage and became almost undetectable until the 8-cell stage (Fig 1B-C). The dynamics of H3K9me3 erasure is rather slow, compared to the rapid erasure of active mark H3K4me3 which occurs within the one-cell stage (Fig EV1A, B), suggesting that active erasure is less likely to reduce H3K9me3 levels during the early cleavage stage. Instead, we reasoned that the passive dilution of H3K9me3 during every DNA replication accounts for the early erasure dynamics of H3K9me3, since medaka blastomeres rapidly divide every 30 minutes until the late morula stage (Iwamatsu, 2004).

**Figure 1.**
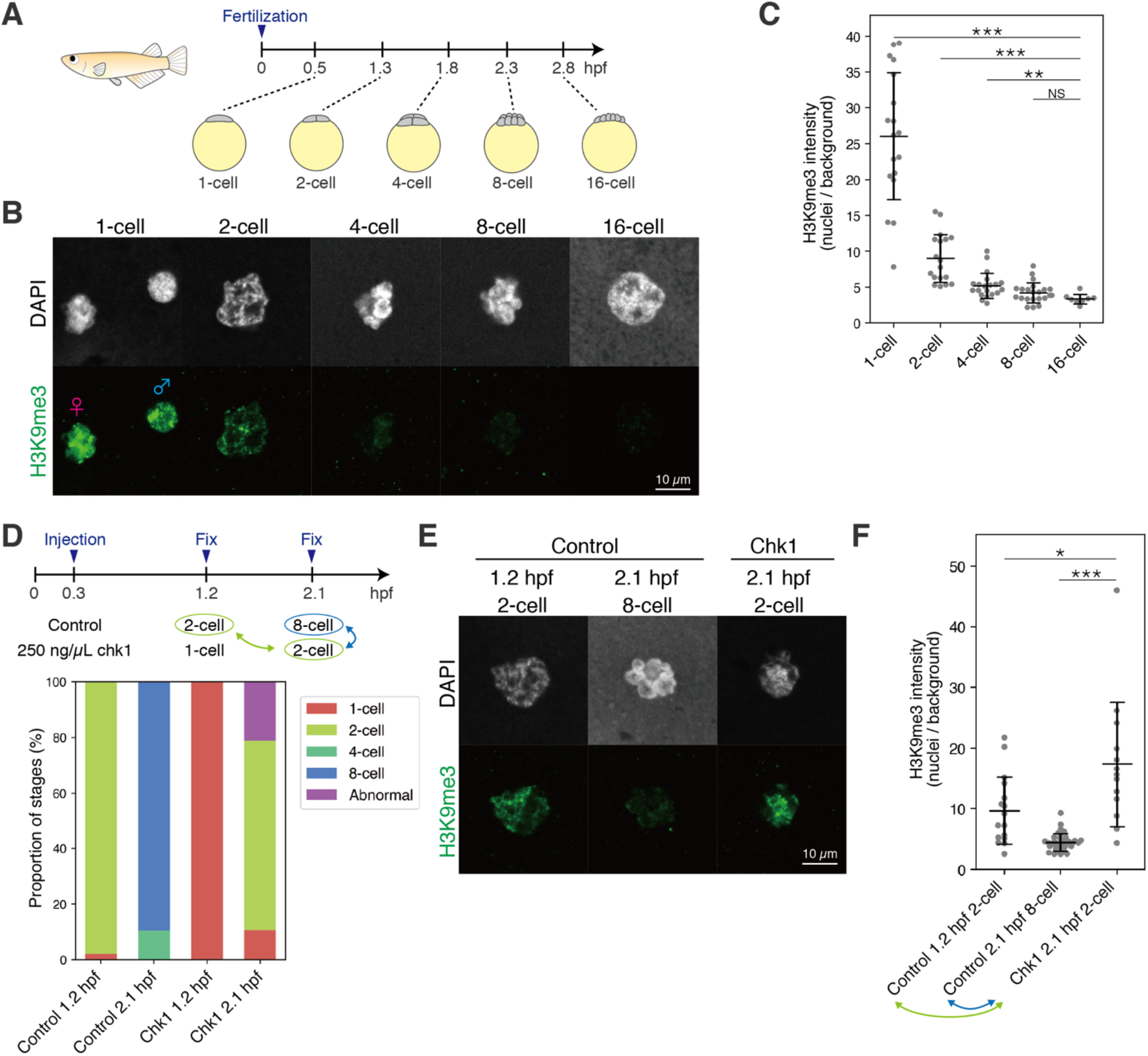
Passive erasure of heterochromatin during early cleavage stages in medaka. (A) Development of medaka embryo during early cleavage stages. (hpf: hours post fertilization) (B) Immunofluorescence staining of H3K9me3 during early cleavage stages. (C) Quantification of (B). Each dot indicates the average intensity of 1-2 cells in a single broad field slice image of single embryo. Two-sided Wilcoxon rank-sum test. Error bars indicate the mean ± s.d. (D) Schematic summarizing the chk1 experiment (top) and the proportion of stages in the chk1 experiment in medaka (bottom). Stages highlighted in green and blue were compared in (E) and (F). (E) Immunofluorescence staining of H3K9me3 in the chk1 experiment. (F) Quantification of (E). Each dot indicates the average intensity of 1-2 cells in a single broad field slice image of single embryo. Two-sided Wilcoxon rank-sum test. Error bars indicate the mean ± s.d *: p < 0.05, **: p< 0.01, ***: p<0.001, NS: not significant.

Thus, we experimentally assessed the dilution hypothesis by prolonging cell cycles. If any active demethylation (e.g. by histone demethylase) functions after fertilization, the prolonged cell cycle should result in a greater reduction of H3K9me3 levels when compared to the control at the same developmental stage. Conversely, if the passive dilution mainly works, H3K9me3 levels should not be affected in embryos with prolonged cell cycles at the same developmental stage. To slow down the cell cycle, we overexpressed Chk1 in cleavage-stage embryos, which inhibits the formation of replication origins (Kappas *et al*, 2000; Chan *et al*, 2019; Collart *et al*, 2017, 2013). In *chk1*-mRNA-injected embryos, the timing of early cleavages was delayed, suggesting the prolonged cell cycle (Fig 1D). In these injected embryos, the level of H3K9me3 detected by immunofluorescence staining was not reduced, while H3K9me3 was almost completely erased in 8-cell stage control embryos, even though the same absolute time after fertilization had elapsed in both embryos (Fig 1D-F, Control 2.1 hpf 8-cell vs Chk1 2.1 hpf 2-cell). Furthermore, prolonged cell cycles did not decrease the H3K9me3 level in injected embryos either, but slightly increased the H3K9me3 level, compared to that in 2-cell stage control embryos (Fig 1D-F, Control 1.2 hpf 2-cell vs Chk1 2.1 hpf 2-cell). Taken together, these data suggest that the erasure of H3K9me3 in the early cleavage stages is mainly caused by DNA-replication dependent dilution in medaka, and that H3K9me3 methyltransferases are active in *de novo* H3K9me3 deposition even during the erasure period (i.e., from the one-cell stage to the 8-cell stage) (discussed later).

### ZGA is dispensable for heterochromatin establishment during the MBT in medaka

In non-mammalian vertebrates, both ZGA and cell cycle slowing take place at the mid-blastula stage, which are collectively called ‘mid-blastula transition’ (MBT) (Tadros & Lipshitz, 2009; Vastenhouw *et al*, 2019). After the global decrease in H3K9me3 at early cleavage stages, medaka embryos begin to re-accumulate H3K9me3 again during the MBT (Fig 2A) (Fukushima *et al*, 2023). Thus, we hypothesized that ZGA and/or the slowing of cell cycle triggers re-accumulation of H3K9me3 during the MBT. First, we investigated the role of ZGA in this process, as previously done in zebrafish (Laue *et al*, 2019). To block ZGA, we injected the RNA polymerase II inhibitorα-amanitin into medaka embryos (Nakamura *et al*, 2021). As expected,α-amanitin injection resulted in developmental arrest and injected embryos did not undergo gastrulation (Fig 2B). The number and morphology of cells were comparable between control and α-amanitin-injected embryos until the late-blastula stage (just before the onset of gastrulation) (Fig 2B, EV2A), indicating that the developmental arrest occurred at the stage of gastrulation. To avoid the effect of developmental arrest, we first compared H3K9me3 levels in control andα-amanitin-injected embryos at the late-blastula stage (the stage just before the developmental arrest becomes evident). As a result, both immunofluorescence staining and quantitative western blot confirmed that the H3K9me3 level was comparable between control andα-amanitin-injected embryos (Fig 2C-F, EV2D, E). The same result was also obtained when compared at the pre-early gastrula stage (beginning of gastrulation) (Fig EV2B, C). We thus conclude that ZGA, or active transcription from the genome, is dispensable for re-accumulation of H3K9me3 during the MBT in medaka.

**Figure 2.**
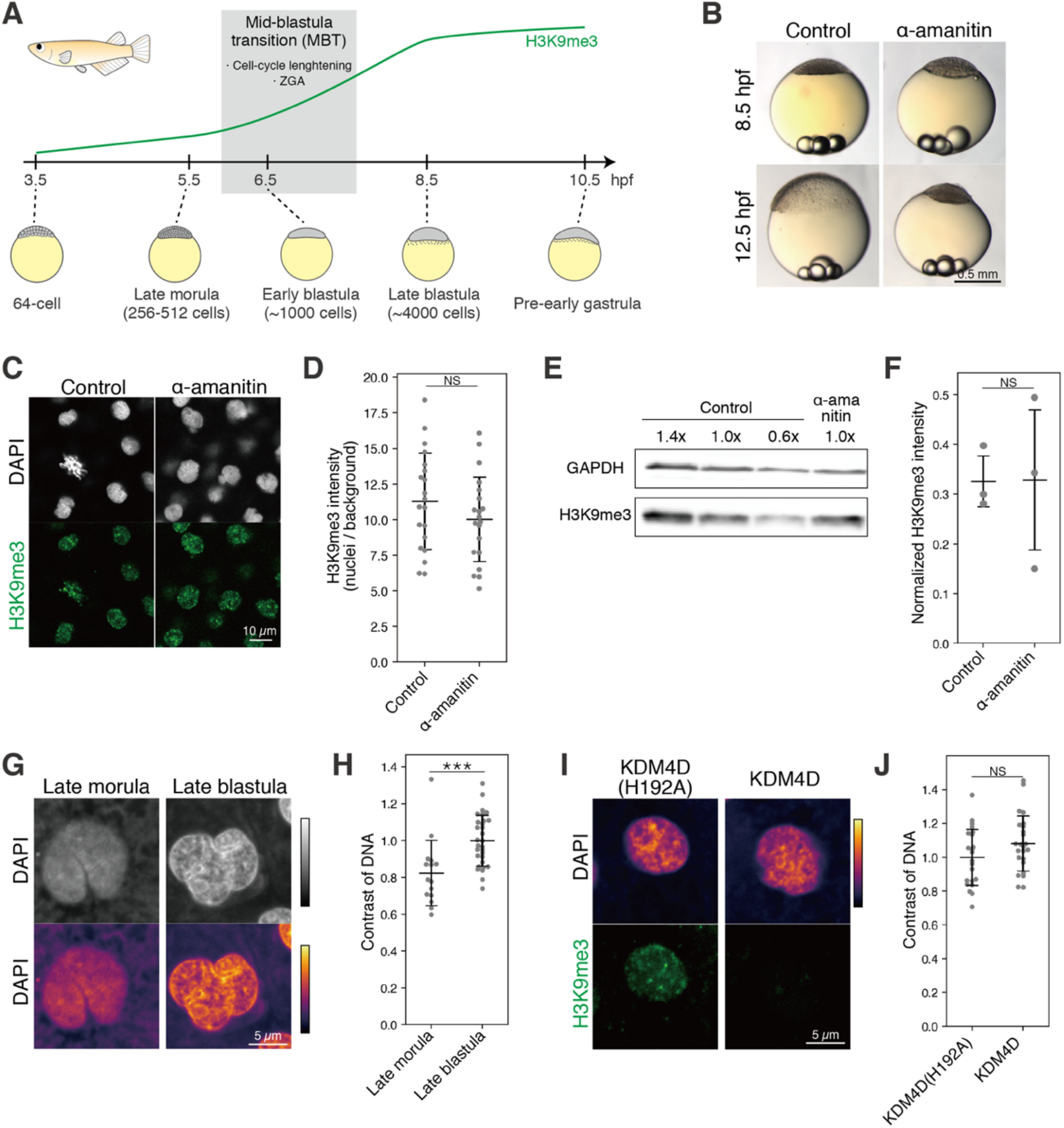
ZGA is dispensable for heterochromatin establishment during the MBT in medaka. (A) Development of medaka embryos before and after the MBT. (B) Phenotype of α-amanitin-injected medaka embryos. (C) Immunofluorescence staining of H3K9me3 in the α-amanitin injection experiment. (D) Quantification of (C). Each dot indicates the average intensity of ∼40 cells in a single broad field slice image of single embryo. Two-sided unpaired Student’s t-test. Error bars indicate the mean ± s.d (E) Western blot of H3K9me3 and GAPDH using control andα-amanitin-injected embryos. (F) Quantification of (E). H3K9me3 signal intensity was normalized by GAPDH signal intensity. Two-sided unpaired Student’s t-test. Error bars indicate the mean ± s.d (G) DAPI-staining at the late morula and late blastula stages. Colormaps are shown at the bottom to better illustrate the appearance of DNA-dense regions at the late blastula stage. (H) Quantification of DNA contrast in (G). Each dot indicates the DNA contrast of a single cell. ∼6 embryos were analyzed. Two-sided Wilcoxon rank-sum test. Error bars indicate the mean ± s.d (I) DAPI and immunofluorescence staining of embryos injected with human KDM4D, a demethylase of H3K9me3, or its catalytically inactive mutant KDM4D(H192A) at the late blastula stage. The pattern of DNA-dense domains was comparable irrespective of presence or absence of H3K9me3. (J) Quantification of DNA contrast in (I). Each dot indicates the DNA contrast of a single cell. Five embryos were analyzed. Two-sided unpaired Student’s t-test. Error bars indicate the mean ± s.d *: p < 0.05, **: p< 0.01, ***: p<0.001, NS: not significant.

### DNA-dense domains detected in nuclei after the MBT do not represent heterochromatin

In the previous studies, in addition to H3K9me3 accumulation, DNA-dense domains in nuclei detected after ZGA was used to indicate the onset of heterochromatin formation (Mutlu *et al*, 2019; Laue *et al*, 2019). We also observed the formation of DAPI-dense domains from the late blastula stage (i.e. after the MBT) in medaka embryos (Fig 2G, H), which is ZGA-dependent as in zebrafish (Laue *et al*, 2019) (Fig EV2F, G). However, a recent study in zebrafish demonstrated that the formation of the DNA-dense domain after the MBT is dependent on micro-phase separation via zygotic transcription (Hilbert *et al*, 2021). Consistently, artificial depletion of H3K9me3 revealed that the formation of DNA-dense domains occurred irrespective of the presence or absence of H3K9me3 (Fig 2I, J). These results favor the idea that DNA-dense domains observed after the MBT do not represent heterochromatin, but probably transcription-dependent condensates (Hilbert *et al*, 2021), and thus cannot be used as an indicator of early-stage heterochromatin formation.

### Cell cycle slowing regulates heterochromatin establishment during MBT in medaka

Since ZGA is not required for H3K9me3 re-accumulation during the MBT, we next assessed if cell cycle slowing regulates this process. Except for earliest cleavages (∼ 30 minutes/round) (Fig 1A)(Iwamatsu, 2004), the cell cycle length during and after the MBT was not precisely determined in medaka. We first counted the cell number per embryo from 3.5 hpf (the 64-cell stage) to 10.5 hpf (the pre-early gastrula stage) (Fig 2A) and found that the cell cycle was initially about 30 minutes/round, gradually prolonged from the 64-cell stage onward, followed by a dramatic slowing from 7.5 hpf, and finally reaching to ∼ 1.7 hours/round towards the late blastula stage (Fig EV3A-C). Thus, there is an apparent correlation between the onset of H3K9me3 re-accumulation and cell cycle slowing in medaka embryos.

To test the causal relationship between the above two events, we prolonged the cell cycle by overexpressing Chk1 at a milder concentration than in the above experiment during the early cleavage stages in Fig 1 (Kappas *et al*, 2000; Chan *et al*, 2019; Collart *et al*, 2017, 2013) and compared H3K9me3 levels by immunofluorescence staining. The developmental delay of *chk1*-injected embryos was evident from the 64-cell stage and onward (Fig 3A), suggesting that the cell cycle was artificially prolonged after the erasure of H3K9me3 (Fig 1A-C). In these *chk1*-injected embryos, we observed an increase in H3K9me3 levels at the morula stage (Fig 3B, C, control 5.5 hpf late morula vs Chk1 8.5 hpf late morula). As an alternative approach, we extended cell cycle length by incubating embryos with the translation inhibitor, cycloheximide (CHX), from the 8-cell stage (i.e., after the erasure of H3K9me3 shown in Fig 1A-C) (Chan *et al*, 2019). This treatment immediately slowed development, and again increased H3K9me3 levels at the 16-cell stage (Fig 3D-F, DMSO 2.8 hpf 16-cell vs CHX 3.5 hpf 16-cell). Since H3K9me3 is normally erased by the 8-cell stage (Fig 1A, B), the observed increase in H3K9me3 in both Chk1 and CHX experiments must not be caused by incomplete erasure of H3K9me3, but by precocious re-accumulation of H3K9me3. These experiments demonstrated that the slowing of the cell cycle is sufficient to induce H3K9me3 re-accumulation at the early stages.

**Figure 3.**
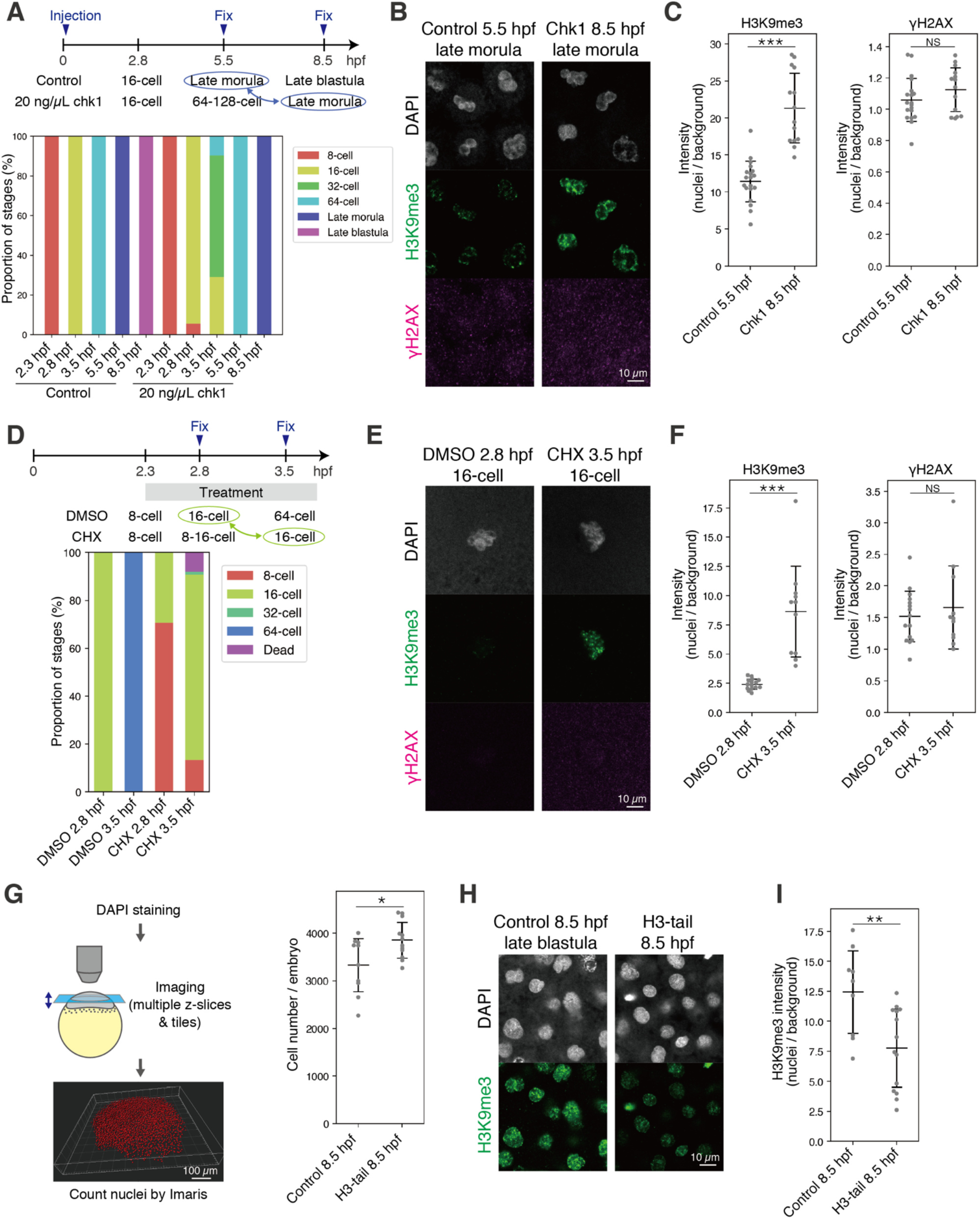
Cell cycle slowing regulates heterochromatin establishment during the MBT in medaka. (A) Schematic summarizing the chk1 experiment (top) and the proportion of stages in the chk1 experiment in medaka (bottom). Stages highlighted in blue were compared in (B) and (C). Error bars indicate the mean ± s.d. (B) Immunofluorescence staining of H3K9me3 and γH2AX in the chk1 injection experiment. (C) Quantification of (B). Each dot indicates the average of 30-40 cells in a single broad field slice image of single embryo. Two-sided Welch’s t-test and two-sided unpaired Student’s t-test were performed for H3K9me3 and γH2AX, respectively. (D) Schematic summarizing CHX treatment (top) and proportion of stages in CHX treatment in medaka (bottom). Stages highlighted in green were compared in (E) and (F). (E) Immunofluorescence staining of H3K9me3 and γH2AX in CHX treatment. (F) Quantification of (E). Each dot indicates the average of ∼5 cells in a single broad field slice image of single embryo. Two-sided Welch’s t-test and two-sided Wilcoxon rank-sum test were performed for H3K9me3 and γH2AX, respectively. Error bars indicate the mean ± s.d. (G) Schematic showing counting of nuclei in an embryo (left) and the number of cells per embryo in the H3-tail injection experiment (right). Two-sided unpaired Student’s t-test. Error bars indicate the mean ± s.d. (H) Immunofluorescence staining of H3K9me3 in H3-tail injection. (I) Quantification of (H). Each dot indicates the average intensity of ∼140 cells in a single broad field slice image of single embryo. Two-sided unpaired Student’s t-test. Error bars indicate the mean ± s.d. *: p < 0.05, **: p< 0.01, ***: p<0.001, NS: not significant.

The above results did not support the ZGA-dependent re-accumulation of H3K9me3, but the possibility still remained that H3K9me3 deposition is dependent on a “timer”, or absolute time elapsed after fertilization. To further test the necessity of cell cycle slowing for H3K9me3 re-accumulation, we accelerated the cell cycle by overexpressing H3-tail, which competitively inhibits Chk1-dependent cell cycle slowing during the MBT (Shindo & Amodeo, 2021). Indeed H3-tail injection accelerated embryonic development at the cleavage stages (Fig EV3D), and the number of cells in H3-tail-injected embryos was higher than that in control at the late blastula stage (Fig 3G), suggesting accelerated cell cycles. Under these conditions, H3K9me3 levels in H3-tail-injected embryos were reduced compared to the control at the late blastula stage, although the same absolute time had elapsed after fertilization (Fig 3H, I). Thus, these data suggest that the slowing of the cell cycle, but not the elapsed absolute time, is critical for the sufficient re-accumulation of H3K9me3 upon the MBT.

The above cell cycle manipulations could induce replication stress or DNA damage which may affect the level of H3K9me3 accumulation. Indeed, it was reported that H3K9me3 depositions were induced by the DNA-repair pathway (Ayrapetov *et al*, 2014). We thus quantified the DNA damage marker, γH2AX, in the cell-cycle manipulation experiments. However, we did not find any statistically significant accumulation of γH2AX in the cell-cycle manipulated embryos (Fig 3B, C, E, F). We also observed that DNA damage, induced by ultraviolet (UV) exposure of late-blastula embryos, increased γH2AX levels but did not H3K9me3 levels (Fig EV3E, F), suggesting that DNA repair-mediated H3K9me3 deposition is less likely to be active in early medaka embryos. Taken together, we conclude that the slowing of the cell cycle causes re-accumulation of H3K9me3 upon the MBT in medaka.

### Setdb1 accumulates in nuclei during the MBT in medaka

To elucidate the molecular mechanism underlying the onset of H3K9me3 deposition during the MBT, we focused on the evolutionary conserved H3K9me3 methyltransferases Suv39h1/2 and Setdb1, which are known to be essential for re-accumulation of H3K9me3 in mouse, and *Drosophila* and *C. elegans*, respectively (Mutlu *et al*, 2018, 2019; Seller *et al*, 2019; Burton *et al*, 2020).

In medaka, Setdb1 and Suv39h1 are encoded by *setdb1b* and *suv39h1b*, respectively. We previously showed that the expression levels of *setdb1b* and *suv39h1b* transcripts are almost comparable before and after the MBT (Fukushima *et al*, 2023). Furthermore, we found that H3K9me3 re-accumulation proceeded even under conditions of translational inhibition by CHX treatment (Fig 3D-F). Thus, neither transcriptional nor translational regulation of *setdb1b* and *suv39h1b* could account for the cell cycle-dependent H3K9me3 deposition during the MBT in medaka.

As another possibility, we focused on post-translational regulation of the histone methyltransferases. Indeed, Setdb1 has both a nuclear export signal (NES) and a nuclear localization signal (NLS) at its N-terminus (Fig EV4A), and the nuclear and cytoplasmic localization of Setdb1 and Eggless (Setdb1 homolog in *Drosophila*) is known to be tightly regulated in mice and *Drosophila*, respectively (Osumi *et al*, 2019; Tachibana *et al*, 2015; Cho *et al*, 2013; Tsusaka *et al*, 2019). Importantly, nuclear accumulation of Eggless and MET-2 (Setdb1 homolog in *C.elegans*) has been implicated in re-accumulation of H3K9 methylations in early development of *Drosophila* and *C. elegans*, respectively (Mutlu *et al*, 2018, 2019; Seller *et al*, 2019). We thus reasoned that nuclear accumulation of maternally provided Setdb1 is associated with the onset of H3K9me3 accumulation during the MBT in medaka, and performed imaging of Setdb1 localization by immunofluorescence staining. After having validated the antibody specificity (Fig EV4A, B), we found that before the MBT, Setdb1 was excluded from the nuclei and mainly localized in the cytoplasm, whereas after the MBT it was localized in both nuclei and cytoplasm (Fig 4A-C). As shown above, the cell cycle length after the MBT becomes much longer than that before the MBT: the late morula stage (5.5 hpf), the late blastula stage (8.5 hpf) and the pre-early gastrula stage (10.5 hpf) is ∼ 0.6, 1.7 and 2.6 hours/round (Fig EV3A-C). This correlation led us to test the causality between cell cycle slowing and the onset of Setdb1 nuclear localization, and we found that cell cycle extension by *chk1* overexpression promoted Setdb1 nuclear localization (Fig 4D). Contrary to Setdb1, Suv39h1 does not have NES (Padeken *et al*, 2022). Consistently, overexpressed FLAG-tagged Setdb1 exclusively localized to the cytoplasm like endogenous one, whereas overexpressed FLAG-tagged Suv39h1 localized to the nuclei at the late morula stage (before the MBT) (Fig EV4C), suggesting that Suv39h1 contributes minimally to the onset of de novo H3K9me3 deposition. These results suggest that, as in *C. elegans* and *Drosophila*, the slowing of the cell cycle induces nuclear accumulation of Setdb1, and thereby leads to re-accumulation of H3K9me3 at the MBT in medaka (Fig 4E).

**Figure 4.**
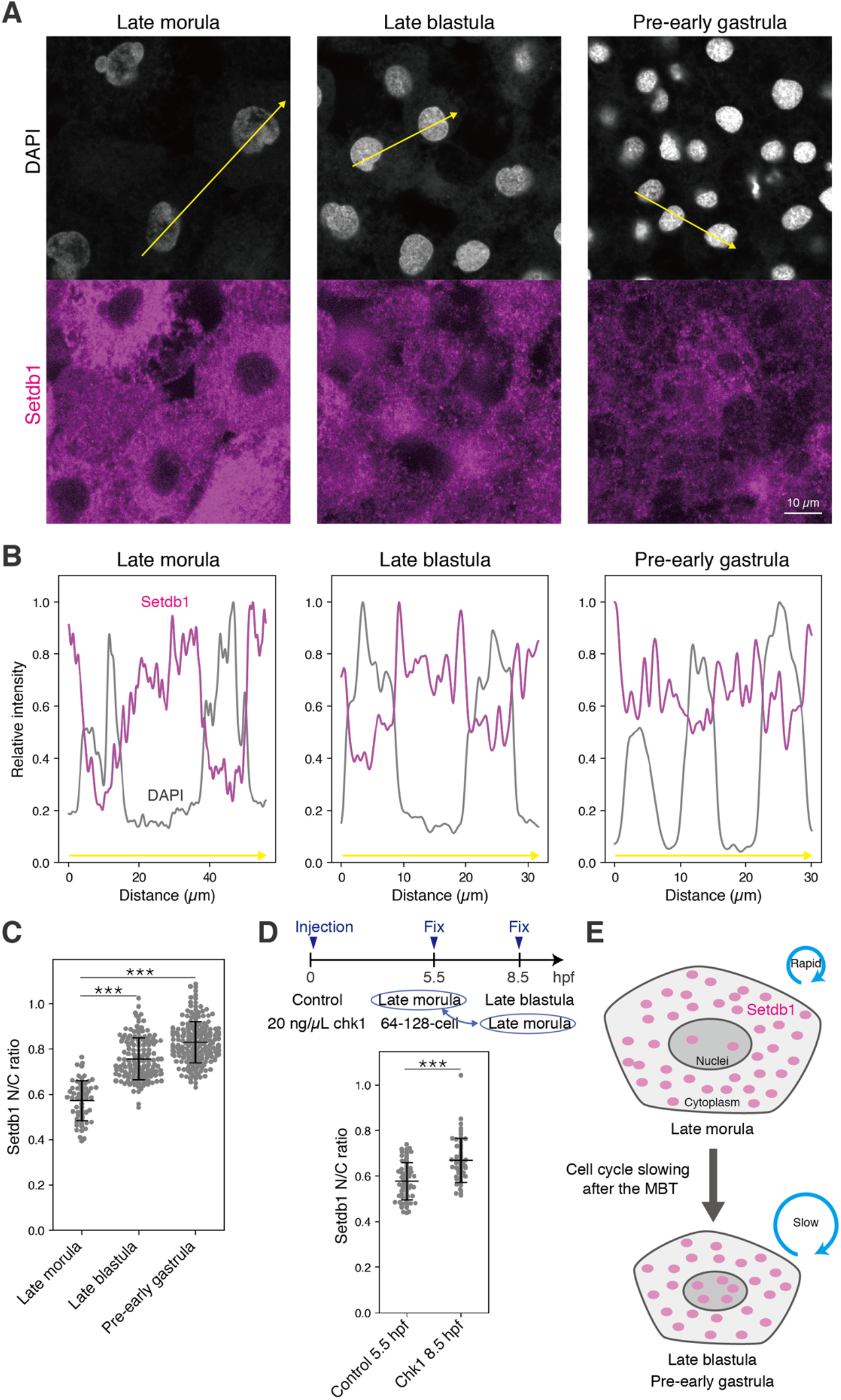
Setdb1 accumulates to nuclei upon the MBT in medaka. (A) Immunofluorescence staining of Setdb1 in medaka embryos before and after the MBT. Signal intensities along the yellow arrows were quantified in (B). (B) Quantification of signal intensity of DAPI and Setdb1 along the yellow arrows in (A). (C) Quantification of nuclear / cytoplasmic ratio (N/C ratio) of Setdb1 in medaka embryos before and after the MBT. Each dot indicates the N/C ratio of a single cell. 10, 8, and 10 embryos at the late morula, late blastula, and pre-early gastrula, respectively, were analyzed. Two-sided Wilcoxon rank-sum test. Error bars indicate the mean ± s.d. (D) Schematic showing chk1 overexpression experiment (top) and quantification of N/C ratio of Setdb1 in control and *chk1*-injected embryos (bottom). Each dot indicates the N/C ratio of a single cell. Eleven embryos were analyzed for each condition. Two-sided Wilcoxon rank-sum test. Error bars indicate the mean ± s.d. (E) Schematic representation of the model of Setdb1 accumulation induced by cell cycle slowing during the MBT. *: p < 0.05, **: p< 0.01, ***: p<0.001, NS: not significant.

### Heterochromatin establishment during the MBT is cell cycle length-dependent in zebrafish

Contrary to our data in medaka, the previous study with zebrafish showed by western blot that H3K9me3 re-accumulation during the MBT depends on ZGA (Laue *et al*, 2019). To address this discrepancy, we first reproduced the previous zebrafish experiment. We first confirmed in zebrafish that H3K9me3 accumulated during the MBT (Fig 5A-C) and then attempted to block ZGA byα-amanitin injection (Laue *et al*, 2019; Chan *et al*, 2019; Zhang *et al*, 2018; Pálfy *et al*, 2020). This resulted in a reduction in the number of cells per embryo at the dome stage (the onset of epiboly) (Fig EV5A, B), and injected embryos did not undergo gastrulation (Fig 5D), indicating that α-amanitin injection leads to developmental arrest at the stage of gastrulation. Under these experimental conditions, we compared H3K9me3 levels between control and α-amanitin-injected embryos at the sphere stage (just before gastrulation) by immunostaining, but found no statistically significant changes (Fig 5E, F). We next compared H3K9me3 levels at the dome stage, which is the stage used in the previous study (Laue *et al*, 2019), but again found no differences (Fig EV5C, D). Our imaging analysis quantified the average signal in nuclei (intensity in nuclei/area of nuclei, see the method for detail) and was not affected by the number of cells per embryo, whereas the previous study (Laue *et al*, 2019) quantified the levels of H3K9me3 by western blot, which is potentially sensitive to the cell number fluctuations. Indeed, our quantitative western blot, in which samples from the same number of embryos at the dome stage were loaded in each lane, showed a reduction in H3K9me3 after α-amanitin injection (Fig EV5E-G) (the similar result as in the previous zebrafish experiment). However, H3K9me3 levels became comparable when normalized by the cell number per embryo based on our data (Fig EV5B, G). Taken together, we conclude that the re-accumulation of H3K9me3 during the MBT is ZGA-independent in zebrafish.

**Figure 5.**
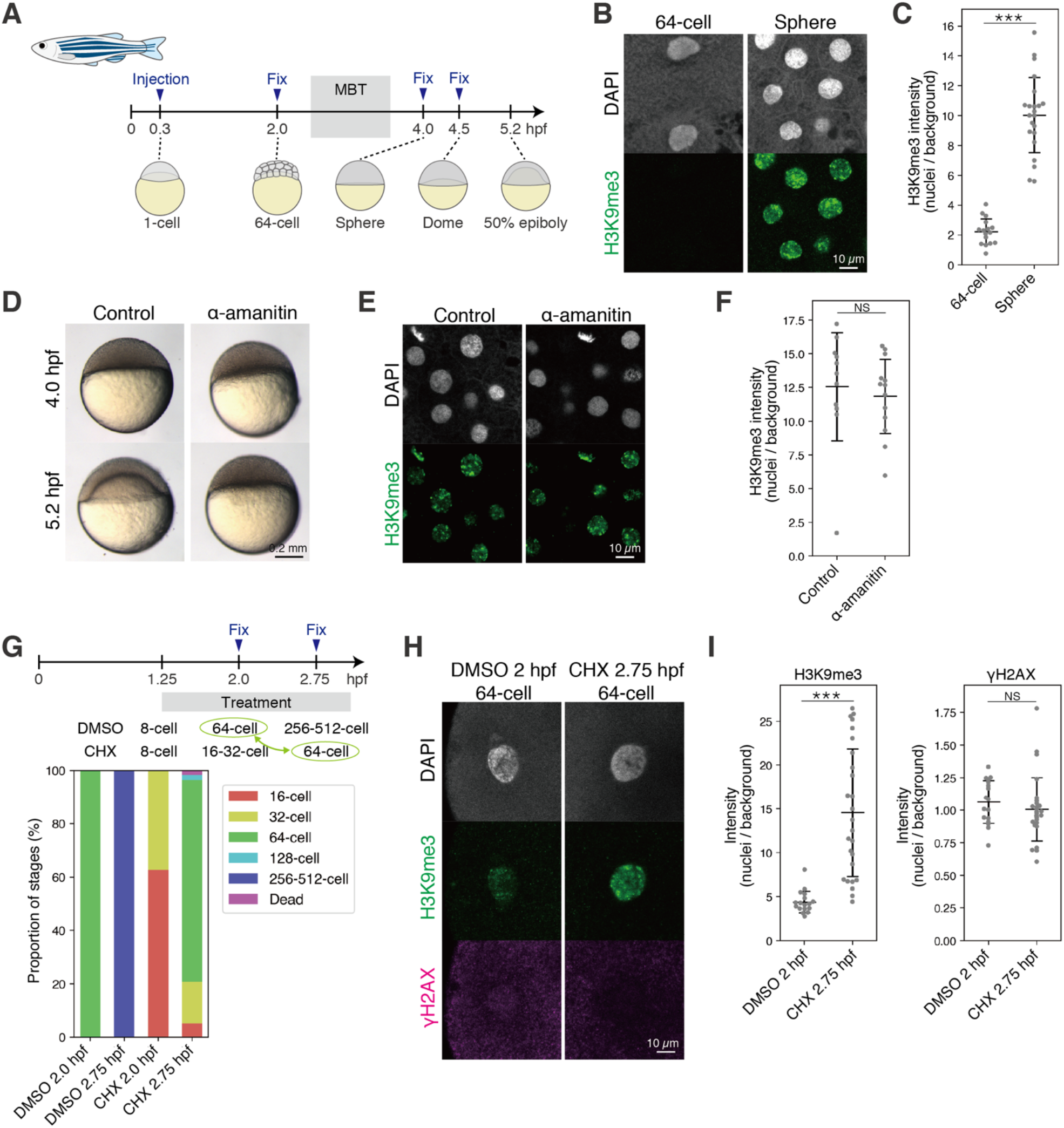
Heterochromatin establishment during the MBT is cell cycle length-dependent in zebrafish. (A) Development of zebrafish embryo before and after the MBT. (B) Immunofluorescence staining of H3K9me3 in zebrafish embryos. (C) Quantification of (B). Each dot indicates the average of ∼5 and ∼60 cells in a single broad field slice image of single embryo at the 64-cell stage and the sphere stage, respectively. Two-sided Welch’s t-test. Error bars indicate the mean ± s.d. (D) Phenotype of α-amanitin-injected zebrafish embryos. (E) Immunofluorescence staining of H3K9me3 in the α-amanitin injection experiment at the sphere stage. (F) Quantification of (C). Each dot indicates the average of ∼60 cells in a single broad field slice image of single embryo. Two-sided Wilcoxon rank-sum test. Error bars indicate the mean ± s.d. (G) Schematic summarizing the CHX experiment (top) and the proportion of stages of CHX-treated zebrafish embryos (bottom). Stages highlighted in green were compared in (H) and (I). (H) Immunofluorescence staining of H3K9me3 and γH2AX in CHX treatment. (I) Quantification of (H). Each dot indicates the average of ∼2 cells in a single broad field slice image of single embryo. Two-sided Wilcoxon rank-sum test. Error bars indicate the mean ± s.d. *: p < 0.05, **: p< 0.01, ***: p<0.001, NS: not significant.

Next, we also tested the cell cycle length dependency of H3K9me3 re-accmulation in zebrafish, and obtained the results similar to those in medaka; CHX treatment from the 8-cell stage prolonged the length of the cell cycle in zebrafish embryos (Fig 5G), and increased H3K9me3 levels at the 16-cell stage without apparent DNA damage (Fig 5H, I). This suggests that the prolonged cell cycle length is sufficient to trigger H3K9me3 deposition in zebrafish. Taken together, the cell cycle length-dependent and ZGA-independent re-accumulation of H3K9me3 is a conserved feature at least among teleosts.

### Heterochromatin establishment during the MBT is cell cycle length-dependent in *Xenopus laevis*

Finally, we extended our analysis to amphibian embryos, *Xenopus laevis*, using immunostaining. As in medaka and zebrafish, H3K9me3 levels gradually increase during the MBT in *Xenopus* (Fig 6A-C). As previously reported (Sudou *et al*, 2016; Chen *et al*, 2019), blocking ZGA by α-amanitin injection resulted in developmental arrest, and injected embryos did not undergo gastrulation (Fig 6D). We tested the requirement of ZGA for H3K9me3 accumulation, but we did not find statistically significant differences in H3K9me3 levels between control and α-amanitin-injected embryos at the stage 10 (the onset of gastrulation) (Fig 6 B, C). We conclude that ZGA is dispensable for H3K9me3 re-accumulation during the MBT also in *Xenopus laevis*.

**Figure 6.**
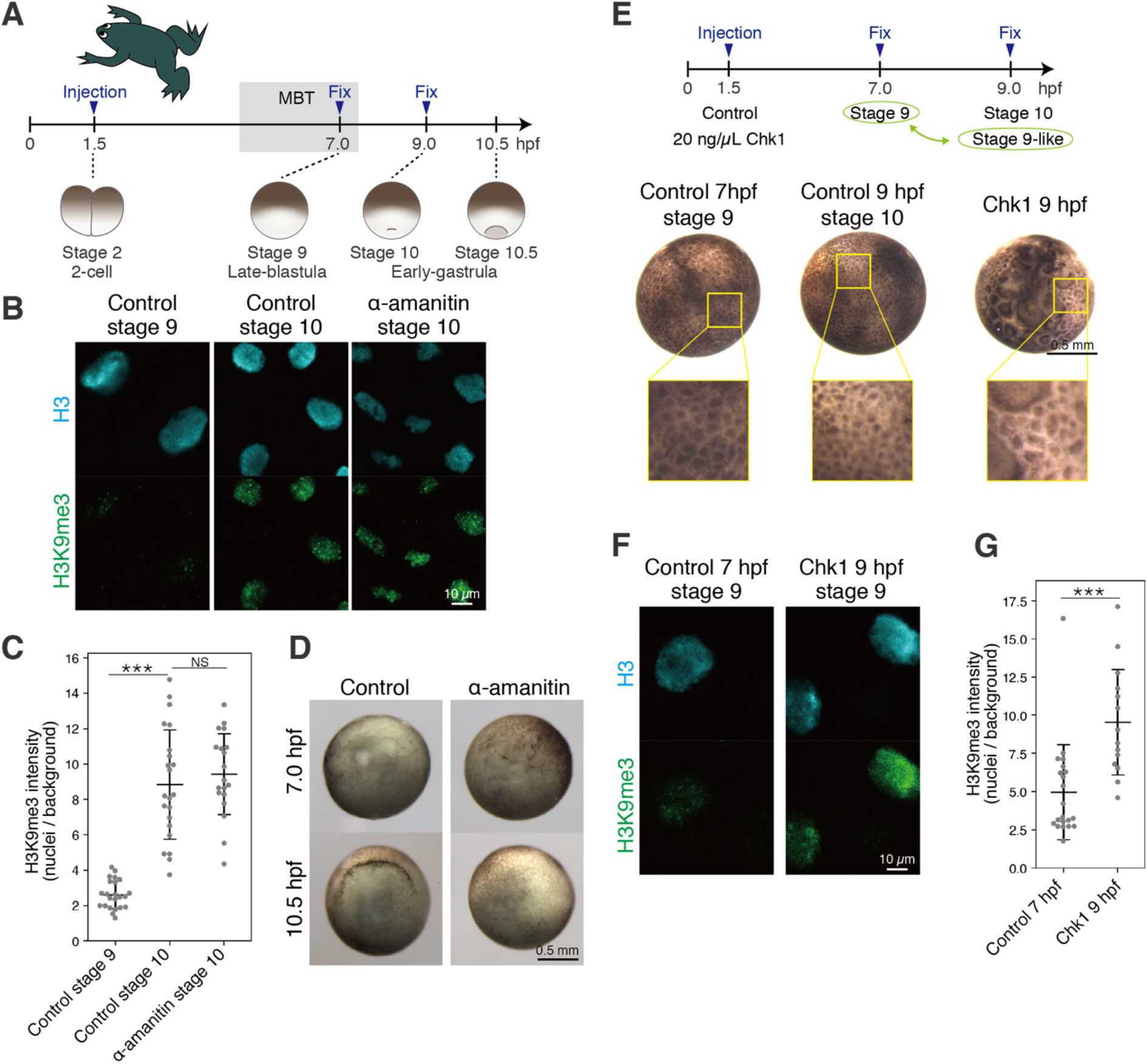
Heterochromatin establishment during the MBT is cell cycle length-dependent in *X. laevis*. (A) Schematic illustration of *X. laevis* embryo development before and after the MBT. (B) Immunofluorescence staining of H3K9me3 in control andα-amanitin-injected *X. laevis* embryo. (C) Quantification of (C). Each dot indicates the average of ∼20 cells in a single broad field slice image of single embryo. Two-sided Welch’s t-test. Error bars indicate the mean ± s.d. (D) Phenotype of α-amanitin-injected *X. laevis* embryos. Unlike control embryos, α-amanitin-injected embryos failed to form dorsal lip at 10.5 hpf, indicating defects in gastrulation. (E) Schematic summarizing *chk1* injection (top) and animal view of *chk1*-injected *X. laevis* embryos (bottom). Stages highlighted in green were compared in (F) and (G). To compare developmental stages, cells at the animal poles are shown in magnified views (yellow squares). (F) Immunofluorescence staining of H3 and H3K9me3 in the *chk1* injection. (G) Quantification of (F). Each dot indicates the average of ∼10 cells in a single broad field slice image of single embryo. Two-sided Wilcoxon rank-sum test. Error bars indicate the mean ± s.d. *: p < 0.05, **: p< 0.01, ***: p<0.001, NS: not significant.

Next, we assessed the cell cycle length dependency by experimentally manipulating the cell cycle length. For this purpose, we first applied *chk1* overexpression in *Xenopus* embryos. The *chk1*-mRNA-injection increased the cell size, although there was variation in size, suggesting that Chk1 overexpression prolonged cell cycles (Fig 6E). For comparison, we selected cells in *chk1*-injected embryos (9 hpf) whose size is almost comparable to that in normal embryos at 7 hpf and found that H3K9me3 levels were higher in *chk1*-injected embryos than that in 7 hpf-normal embryos (Fig 6F, G), indicating that cell cycle slowing caused precocious accumulation of H3K9me3. The same result was obtained by CHX treatment from the 64-cell stage; prolonged cell cycles (Fig EV6A) caused precocious accumulation of H3K9me3 (Fig EV6A-C). Collectively, the mechanism of H3K9me3 re-accumulation during the MBT in *Xenopus* is likely to be similar to that in teleosts, i.e. ZGA-independent and cell cycle length-dependent.

## Discussion

The proper reboot of the epigenetic memory after fertilization is an essential process for early embryogenesis, but its molecular mechanisms were not fully understood in vertebrates. In this study, we have shown which factor mediates heterochromatin erasure and re-establishment during early development in medaka, zebrafish, and *Xenopus*. We first showed that the erasure of H3K9me3 was caused by DNA replication-dependent passive dilution. Second, we revealed that it is cell cycle slowing, not ZGA, that triggers the re-accmulation of H3K9me3 during the MBT. When rapid embryonic cell cycles slow down toward the MBT, nuclear localization of maternal Setdb1 and accumulation of H3K9me3 simultaneously became detectable. This led us to propose the model that Setdb1 accumulated in nuclei during prolonged cell cycles promotes the deposition of H3K9me3. In other words, rapid cell cycles limit nuclear accumulation of Setdb1, but prolonged cell cycle permits accumulation of Setdb1, leading to the onset of H3K9m3 deposition. Consistent with this model, experimental prolongation of the cell cycle increased H3K9me3 levels at the early cleavage stages (Fig 1D-F). The data obtained in zebrafish, medaka and *Xenopus* are all consistent with each other. Our study thus provides experimental evidence for the essential role of cell cycle length in both the erasure and re-establishment of heterochromatin during early development in non-mammalian vertebrates (Fig 7).

**Figure 7.**
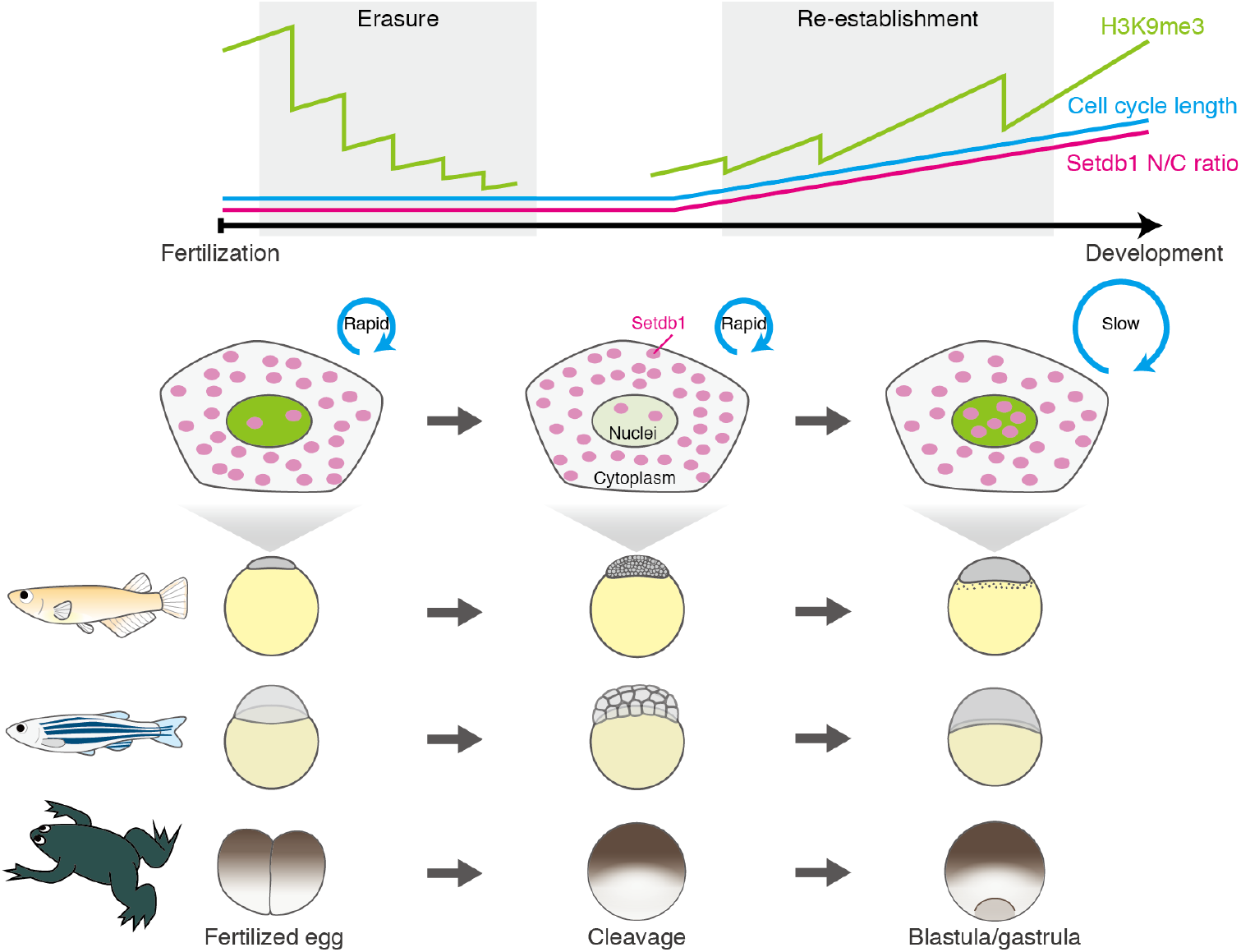
Cell cycle length regulates both erasure and re-establishment of heterochromatin during early development in non-mammalian vertebrates. Schematic summarizing the model of H3K9me3 reprogramming in non-mammalian vertebrates. Cell cycles in fertilized eggs and cleavage embryos are very rapid in non-mammalian vertebrates. This prevents Setdb1 (magenta dots) from accumulating in nuclei, resulting in DNA replication-dependent gradual erasure of H3K9me3 (green intensity in nuclei). However, cell cycles were prolonged from the MBT. Hereafter, Setdb1 can sufficiently accumulate in nuclei during the slowing of cell cycles, increasing H3K9me3 levels in blastula and gastrula embryos.

We showed that ZGA is dispensable for the onset of H3K9me3 deposition in non-mammalian vertebrates. In contrast, it was previously reported in zebrafish that zygotic transcription of miR-430 and its mediated degradation of maternal factors are essential for this process (Laue *et al*, 2019). As described in Results, this discrepancy can be resolved by taking into account developmental arrest and the resulting reduction in cell number. However, we do not exclude the possibility that the degradation of maternal factors is also involved in the re-accumulation of H3K9me3.

Similar to non-mammalian vertebrates, invertebrate embryos such as *Drosophila* and *C. elegans* undergo rapid cleavages in early development (Tadros & Lipshitz, 2009; Vastenhouw *et al*, 2019) and re-install H3K9 methylation after 5-10 rounds of cleavages (Mutlu *et al*, 2018, 2019; Seller *et al*, 2019). In these species, cell cycle slowing also times heterochromatin formation, which is mediated by translocation of Setdb1 homolog from the cytoplasm to nuclei (Mutlu *et al*, 2018, 2019; Seller *et al*, 2019). Therefore, the molecular mechanism of H3K9me3 re-accumulation during epigenetic reprogramming is broadly conserved in rapid-cleavage species. During rapid cleavage, transcription machinery is robustly inactivated before ZGA, due to the absence of “activators” (Jukam *et al*, 2017), lack of active histone modifications (Chan *et al*, 2019), excessive nucleosome (Joseph *et al*, 2017; Amodeo *et al*, 2015; Wilky *et al*, 2019), insufficient nuclear import machinery (Shen *et al*, 2022) and so on. Under these conditions, heterochromatin-mediated gene silencing is not necessarily required. Furthermore, active demethylation is also not required (Mutlu *et al*, 2019), as passive dilution is sufficient to erase H3K9me3 globally and almost completely by the onset of ZGA (after several rounds of cell division). The only exception reported so far is the retention of H3K9me3 at the telomeric region, which is essential for the maintenance of genomic stability (Fukushima *et al*, 2023). This specific retention can be interpreted in the context of the passive erasure, if we assume that the original level of H3K9me3 accumulation is high enough prior to reprograming. Taken together, the passive erasure, which is characterized by a uniform reduction of H3K9me levels along the entire genomic region, would be more cost-effective in rapid-cleavage species.

By contrast, in mammalian embryos, postfertilization cleavages are much slower, and ZGA is initiated in a few rounds of cleavage (Tadros & Lipshitz, 2009; Vastenhouw *et al*, 2019). In this case, the chance of passive erasure is limited, and instead, active demethylation comes into play during the cleavage stages (Sankar *et al*, 2020; Liu *et al*, 2018). This active erasure can work site-specifically and as a result, in mammals H3K9me3 undergo reprogramming in a non-uniform manner (Wang *et al*, 2018; Yu *et al*, 2022; Zhou *et al*, 2023). Mammals take advantage of this mechanism to ensure H3K9me retention in specific genomic regions where it is needed, such as silencing of LTRs (Wang *et al*, 2018) and maintenance of DNA methylation as CpG-rich loci during cleavage stages (Yang *et al*, 2022). In addition, *de novo* H3K9me3 deposition begins immediately after fertilization using Suv39h1/2 (Burton *et al*, 2020). Therefore, we propose a scenario of stepwise evolution of H3K9me3 reprogramming dynamics; the cell cycle-dependent H3K9me3 reprogramming observed in rapid-cleavage species evolved first, and mammals later adopted active demethylation to adjust slow cleavage and immediate ZGA.

**Figure EV1.**
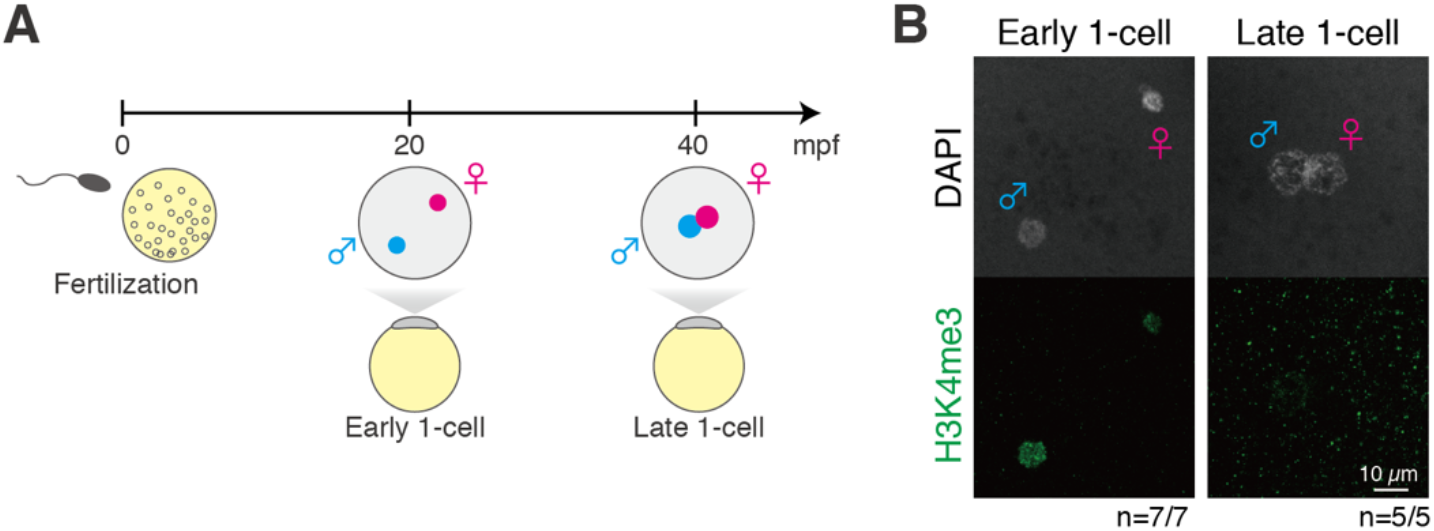
Supportive data for Fig 1. (A) Development of medaka embryos at the one-cell stage. Blue and magenta indicate paternal and maternal pronuclei, respectively. (B) Immunofluorescence staining of H3K4me3 at the one-cell stage. The number of embryos with the representative pattern was indicated at the bottom.

**Figure EV2.**
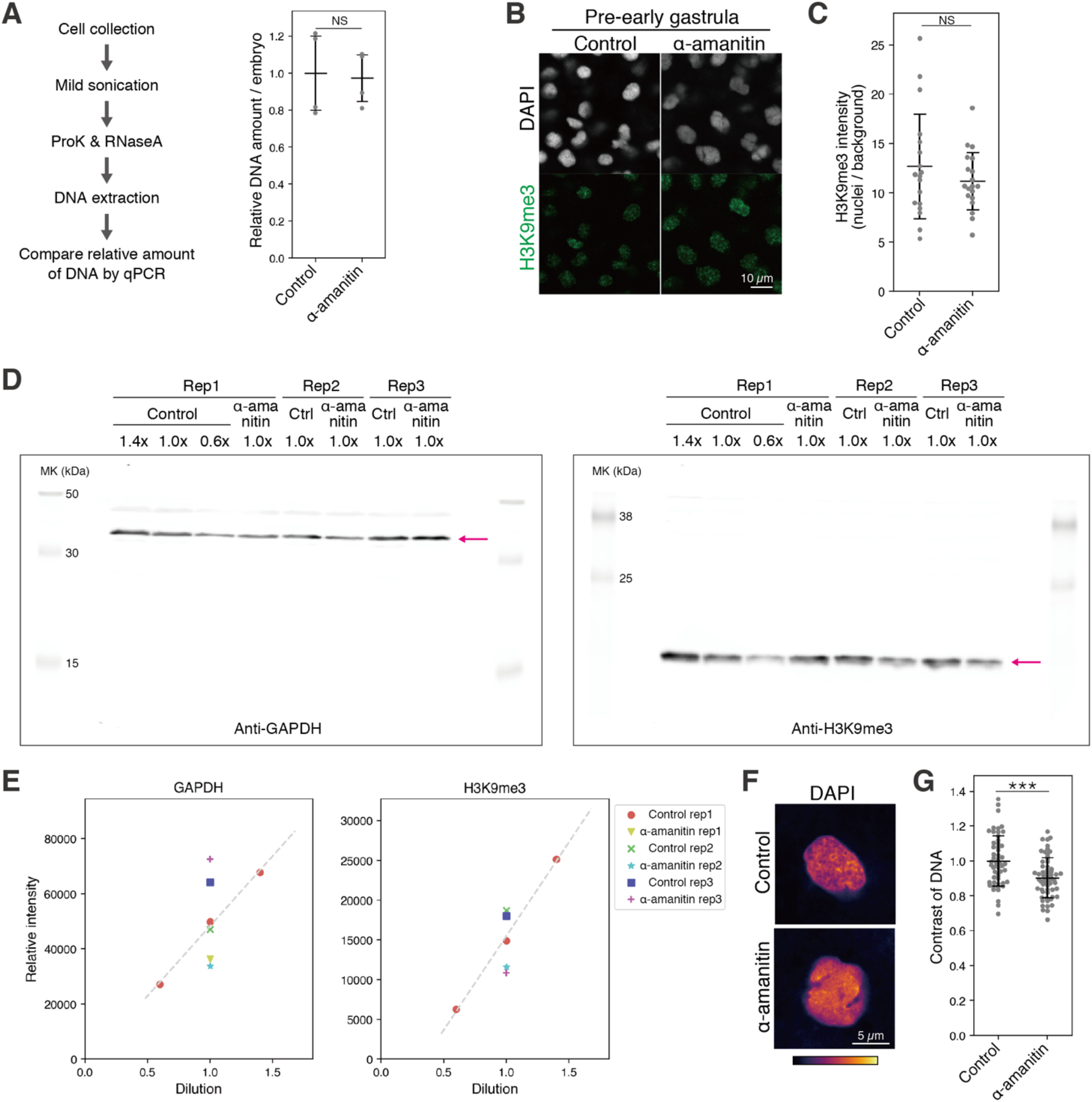
Supportive data for Fig 2. (A) Procedure of quantification of relative amount of DNA per embryo (left) and the results (right) at the late blastula stage. Two-sided unpaired Student’s t-test. Error bars indicate the mean ± s.d. (B) Immunofluorescence staining of H3K9me3 inα-amanitin-injected medaka embryos at the pre-early gastrula stage. (C) Quantification of (B). Each dot indicates the average intensity of ∼50 cells in a single broad field slice image of single embryo. Two-sided Welch’s t-test. Error bars indicate the mean ± s.d. (D) Uncropped results of quantitative western blot using anti-GAPDH and anti-H3K9me3 antibodies. Magenta arrows indicate the specific bands. (E) Quantification of western blot signal intensities in Fig 1F and EV2D. Scatter plots show that all signal intensities of western blots were within the linear range. (F) DAPI staining of control or α-amanitin injected embryos at the late blastula stage. (G) Quantification of (H). Each dot indicates the DNA contrast of a single cell. Ten embryos were analyzed. Two-sided unpaired Student’s t-test. Error bars indicate the mean ± s.d. *: p < 0.05, **: p< 0.01, ***: p<0.001, NS: not significant.

**Figure EV3.**
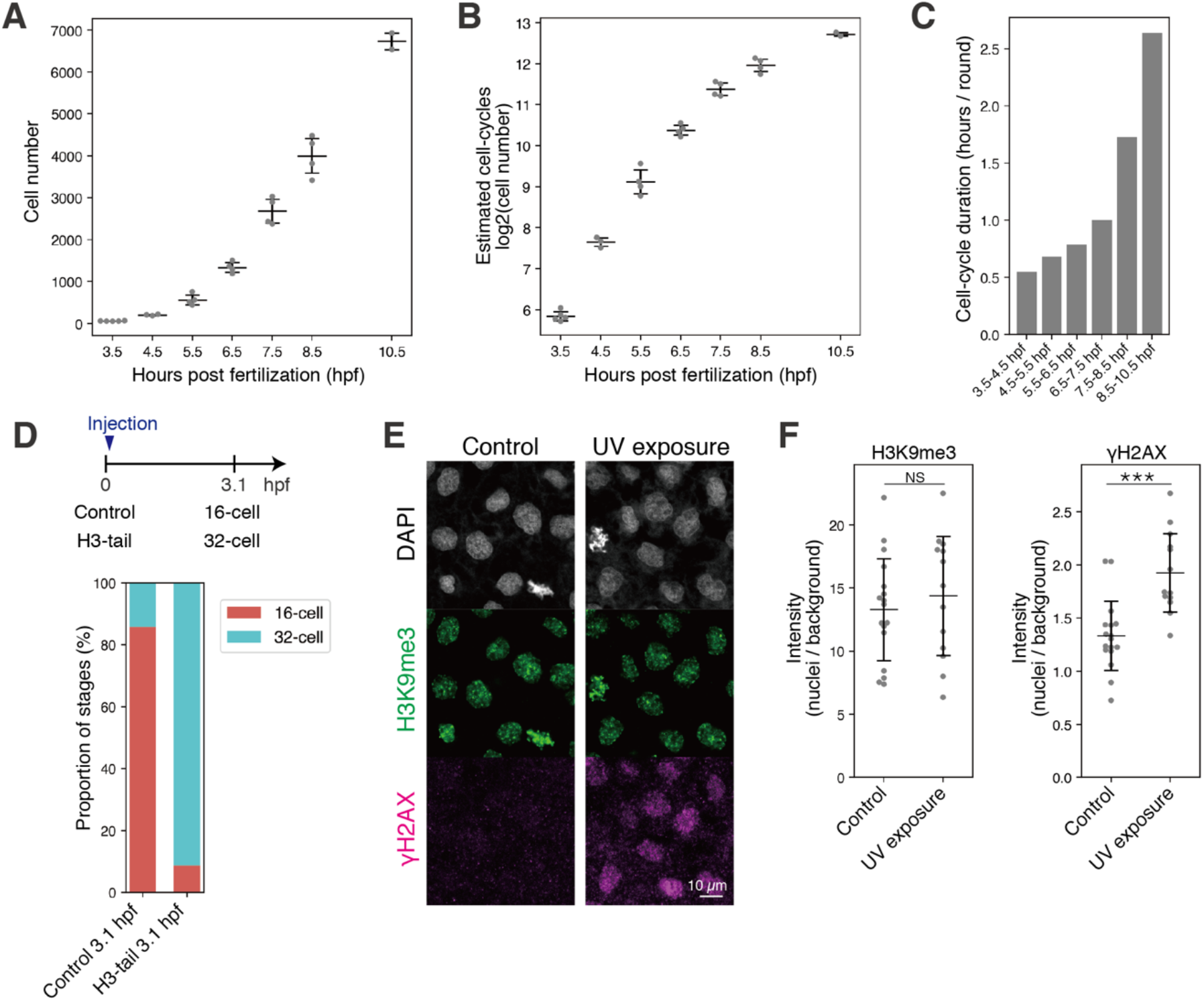
Supportive data for Fig 3. (A) Number of cells per embryo at 3.5-10.5 hpf counted by Imaris software using DAPI staining data. Error bars indicate the mean ± s.d. (B) Number of post fertilization cell-cycles estimated by total number of cells per embryo in (A). Error bars indicate the mean ± s.d. (C) Cell cycle length estimated by cell cycle number and time line in (B). (D) Schematic summarizing H3-tail injection (top) and proportion of stages of H3-tail-injected embryos in the cleavage stages (bottom). (E) Immunofluorescence staining of H3K9me3 and γH2AX in control and UV-treated embryos. (F) Quantification of (B). Each dot indicates the average of ∼100 cells in a single broad field slice image of single embryo. Two-sided unpaired Student’s t-test. Error bars indicate the mean ± s.d. *: p < 0.05, **: p< 0.01, ***: p<0.001, NS: not significant.

**Figure EV4.**
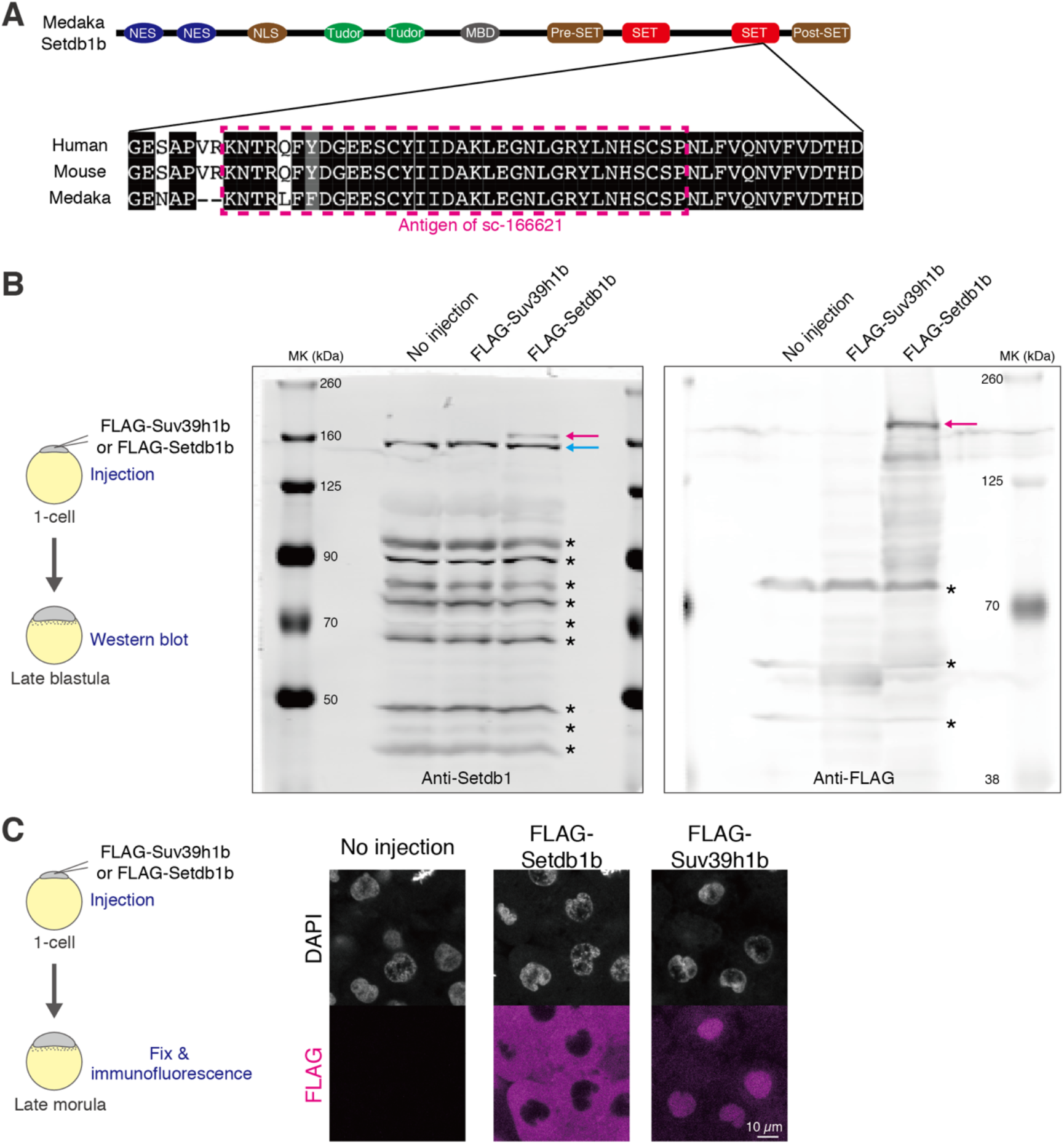
Supportive data for Fig 4. (A) Domains of Setdb1 and amino acid sequence in the SET domain. The antigen sequence of the anti-Setdb1 antibody (sc-166621) is highlighted in magenta. (B) Schematic of the experiments to validate the specificity of the anti-Setdb1 antibody (sc-166621) (left) and the results of western blot (right). Blue and magenta arrows indicate endogenous Setdb1 and exogenously expressed FLAG-Setdb1, respectively. Asterisks (*) indicate non-specific bands. (C) Schematic of the experiments to validate the localization of exogenously expressed FLAG-Suv39h1 and FLAG-Setdb1 (left) and immunofluorescence staining against anti-FLAG at the late morula stage (right). Consistent with the Fig 4, exogenously overexpressed FLAG-Setdb1 localized to cytoplasm, while FLAG-Suv39h1 mainly accumulated in nuclei.

**Figure EV5.**
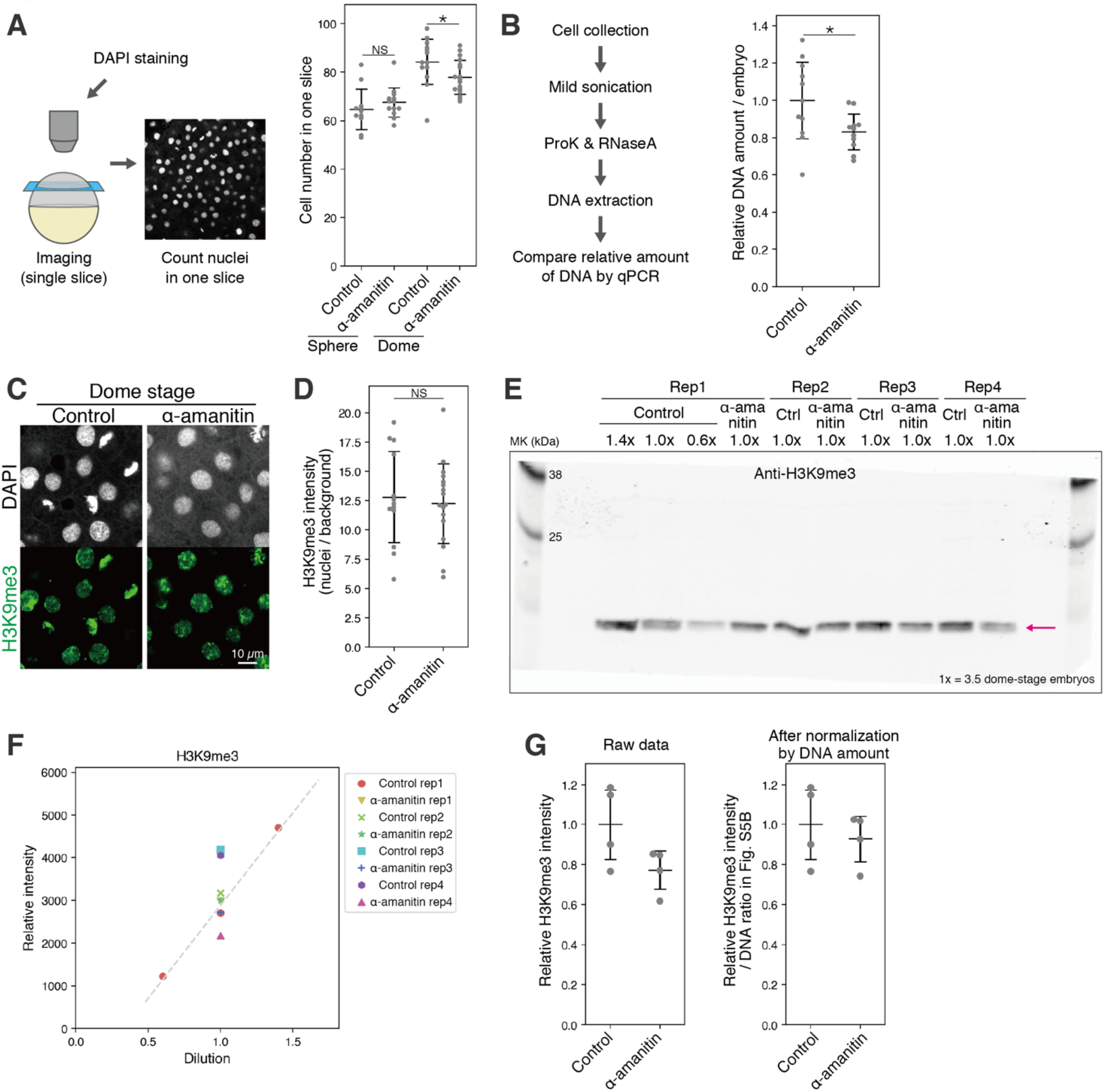
Supportive data for Fig 5. (A) Schematic of counting nuclei in an embryo using a single slice (left) and the cell number in α-amanitin-injected zebrafish embryos (right). Two-sided unpaired Student’s t-test. Error bars indicate the mean ± s.d. (B) Procedure of quantification of the relative amount of DNA per embryo (left) and the results (right) in α-amanitin-injected zebrafish embryos at the dome stage. Two-sided Welch’s t-test. Error bars indicate the mean ± s.d. (C) Immunofluorescence staining of H3K9me3 in α-amanitin injection experiment at the dome stage. (D) Quantification of (C). Each dot indicates the average of ∼80 cells in a single broad field slice image of single embryo. Two-sided unpaired Student’s t-test. Error bars indicate the mean ± s.d. (E) Uncropped results of quantitative western blot using anti-H3K9me3 antibody. The Magenta arrow indicates the specific bands. The same number of dome-stage embryos (1x = ∼3.5 embryos / lane) were loaded into each lane to compare total H3K9me3 levels per embryo. (F) Quantification of western blot signal intensities in Fig EV5E. Scatter plots shows that all western blot signal intensities were within the linear range. (G) Quantification of (F). On the right, data after normalization by the DNA ratio measured in Fig EV5B. Error bars indicate the mean ± s.d. *: p < 0.05, **: p< 0.01, ***: p<0.001, NS: not significant.

**Figure EV6.**
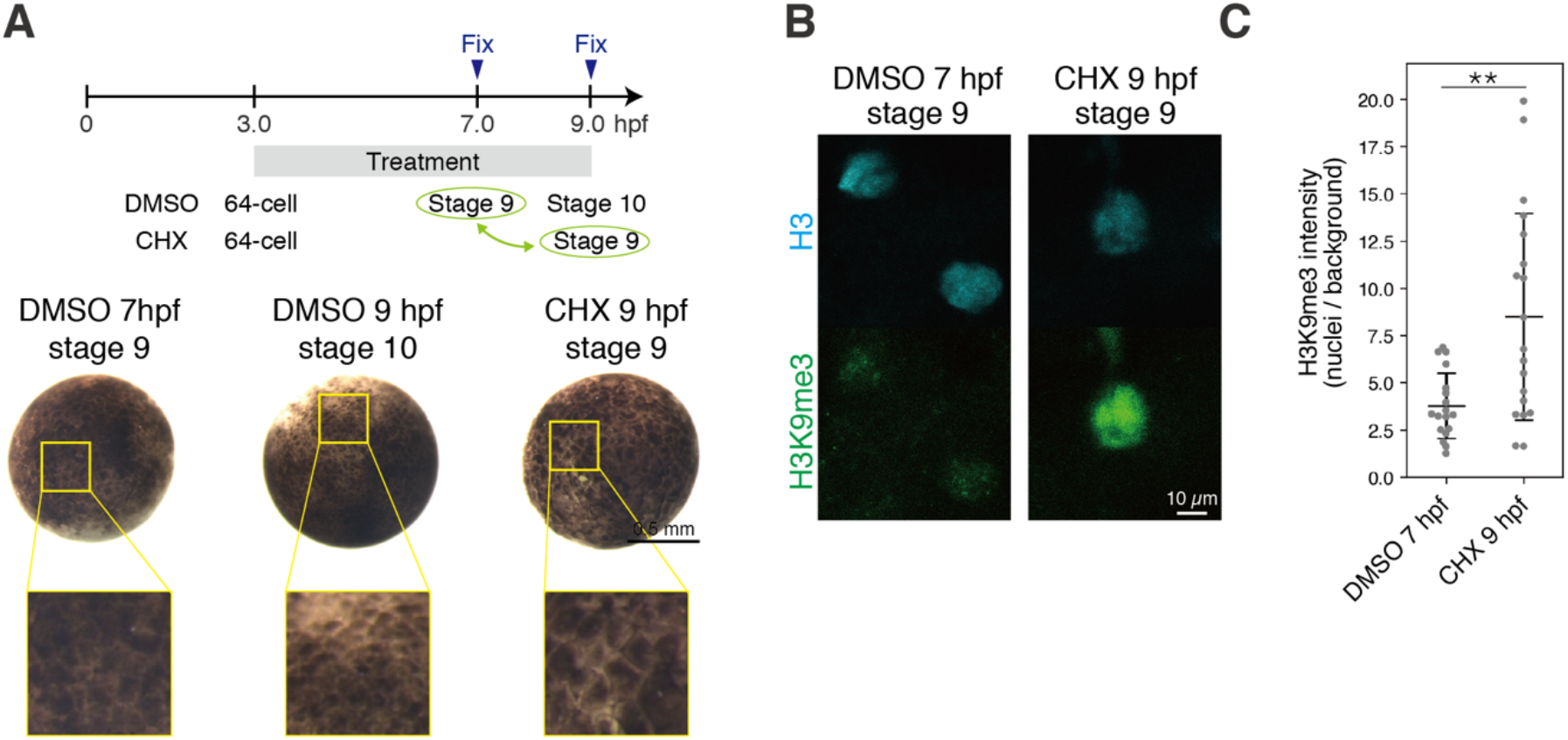
Supportive data for Fig 6. (A) Schematic summarizing the CHX treatment (top) and animal view of CHX-treated *X. laevis* embryos (bottom). Stages highlighted in green were compared in (B) and (C). To compare developmental stages, cells at the animal poles are magnified (yellow squares). (B) Immunofluorescence staining of H3 and H3K9me3 in CHX treatment. (C) Quantification of (F). Each dot indicates the average of ∼5-10 cells in a single broad field slice image of single embryo. Two-sided Welch’s t-test. Error bars indicate the mean ± s.d. *: p < 0.05, **: p< 0.01, ***: p<0.001, NS: not significant.

## Methods

### Animal procedures

Medaka d-rR strain was used in this study. Fertilized embryos were raised according to standard protocols (Kinoshita et al. 2009) at 28℃. Developmental stages were determined according to previously published guidelines (Iwamatsu 2004) (Fig 1A, 2A and EV1A). 10 ng/µLα-amanitin (Nakamura *et al*, 2021), 200 ng/µL *H3-tail* mRNA, 250 or 20 ng/µL medaka’s *chk1* mRNA, 131 ng/µL (300 nM) *FLAG-suv39h1b* mRNA, 358 ng/µL (300 nM) *FLAG-setdb1b* mRNA, 514 ng/µL human *KDM4D* mRNA, and 514 ng/µL human *KDM4D(H192A)* mRNA (Fukushima *et al*, 2023) were injected into one-cell stage embryos. Dechorionated embryos were incubated with 20 ng/µL CHX from the 8-cell stage.

Zebrafish RW strain was used in this study. Fertilized embryos were raised according to standard protocols (Kimmel *et al*, 1995) at 28.5℃. Developmental stages were determined according to previously published guidelines (Kimmel *et al*, 1995) (Fig 5A). One-cell stage embryos were injected with 100pg α-amanitin (Laue *et al*, 2019; Chan *et al*, 2019; Zhang *et al*, 2018; Pálfy *et al*, 2020). Dechorionated embryos were incubated with 20 ng/µL CHX from the 8-cell stage.

Wild-type *X. laevis* was used in this study. *In vitro* fertilized embryos were raised at 23℃. Developmental stages were determined according to previously published guidelines (Nieuwkoop & Faber, 1994) (Fig 6A). At the two-cell stage, 100pg α-amanitin (Sudou *et al*, 2016; Chen *et al*, 2019) and 200 pg medaka *chk1* mRNA were injected into each blastomere. Embryos were incubated with 20 ng/µL CHX from the 64-cell stage.

All experimental procedures and animal cares were performed under the approval of the animal ethics committee of the University of Tokyo (Approval No. 20-2).

### Constructions and *in vitro* transcription

Medaka *chk1*, *setdb1b*, and *suv39h1b* sequences were amplified by PCR from the medaka cDNA library, cloned into TOPO vector using TOPO TA Cloning Kit Dual Promoter (Invitrogen, 45-0640), and introduced to pCS2+ vector using NEBuilder HiFi DNA Assembly Master Mix (NEB, E2621). The H3-tail sequence (Shindo & Amodeo, 2021) was generated by PCR using human histone H3.1 sequence as a template and inserted directly into pCS2+ vector using NEBuilder HiFi DNA Assembly Master Mix. Templates for *in vitro* transcription were amplified by PCR, and mRNA was transcribed *in vitro* from the templates using HiScribe T7 ARCA mRNA Kit with Tailing (NEB, E2060) or mMESSAGE mMACHINE SP6 Transcription Kit (Thermo, AM1340). Previously generated plasmids (Fukushima *et al*, 2023) were used for *in vitro* transcription of human *KDM4D and KDM4D(H192A)* mRNA. All primer sequences used for construction are listed in Table EV1. The transcribed mRNA was purified using RNeasy mini kit (QIAGEN).

### Immunofluorescence staining and imaging

Immunofluorescence staining was performed according to the previous protocol (Fukushima *et al*, 2023) with minor modifications. Briefly, embryos fixed with 4% PFA/PBS were permeabilized with 0.5% Triton X-100/PBS for 15 minutes at room temperature, washed with PBS, incubated with blocking buffer (2% BSA, 1% DMSO, 0.2% Triton X-100, 1×PBS) for 1 hour at room temperature, incubated with primary antibodies overnight at 4°C, washed with PBSDT (1×PBS, 1% DMSO, 0.1% Triton X-100), incubated with blocking buffer for 1 hour at room temperature, incubated with secondary antibodies and DAPI for 4 hours at 4°C, and washed with PBSDT. For immunofluorescence staining of endogenous Setdb1, after permeabilization, samples were incubated with 4 M HCl for 15 minutes at room temperature and incubated with 100 mM Tris-HCl for 20 minutes at room temperature for antigen retrieval. For immunofluorescence staining of medaka embryos, blastodiscs were manually separated from yolk and mounted on slide glasses with coverslips. For immunofluorescence staining of *X. laevis* embryos, whole embryos were mounted on slide glasses with coverslips. For immunofluorescence staining of zebrafish embryos, samples were mounted in 1% low-melting point agarose / PBS. Imaging was performed with a Zeiss LSM710. Total cell count images are z-stacked and tiled images, and the other images are single-slice images. Antibodies used for immunofluorescence are listed in Table EV2.

### UV exposure

8.25 hpf medaka embryos were placed in clean bench and exposed UV for 10 minutes, and incubated for 5 minutes at 28℃. The UV-exposed embryos were then fixed, and immunofluorescence staining was performed as described above.

### Quantification of imaging

Images were quantified using Fiji (Schindelin *et al*, 2012) according to a previous protocol (Fukushima *et al*, 2023) with minor modifications. For medaka and zebrafish samples, DAPI-dense regions were automatically or manually selected as nuclei, signal intensities of histone modifications in nuclei were measured, and the averages were normalized by signal intensities in manually selected background areas. Because the DAPI signal was relatively weak and noisy in *X. laevis* samples, H3-dense regions were automatically or manually selected as nuclei, and signals were measured as above. To measure the N/C ratio of Setdb1, DAPI-dense regions and surrounding regions were manually selected, signal intensities and area sizes were measured, and the N/C ratio was calculated.

### Cell number estimation by qPCR

The same number of dechorionated embryos were homogenized in microtubes by gentile pipetting up and down in ice-cold PBS. After centrifugation, the supernatant was removed, and the pellet was stored in a freezer. The cell pellet was lysed with lysis buffer (50 mM Tris-HCl pH 8.0, 10 mM EDTA, 1% SDS) and sonicated briefly with Covaris (peak power: 105, duty factor: 4.0, cycles per burst: 200, duration: 180 seconds). After RNaseA treatment and ProK treatment, DNA was purified by phenol: chloroform: isoamyl alcohol method and ethanol purification. The relative amount of DNA was measured by qPCR using AriaMx (Agilent). Primers used in this study are listed in Table EV1.

### Total cell number counting

We counted the total number of cells per medaka embryo from images following the protocol of a previous study (Joseph *et al*, 2017). Briefly, tile-scan and z-stack DAPI images were acquired using immunofluorescence staining samples, the figures were stitched using ZEN 2.3 SP1 (ZEISS), and the number of nuclei was counted using Imaris 8.1.2 (BITPLANE).

### Counting cell number in a single slice

Because the blastomere layers of zebrafish blastula were thicker than those of medaka blastula, it is difficult to detect DAPI signals from blastomeres located deep in zebrafish embryos. Therefore, we could not count the total cell number of zebrafish embryos using our systems. To compare total cell number per embryo in parallel with its estimation by qPCR, we counted the number of nuclei in single slice images obtained for immunofluorescence staining (Fig EV5A).

### Estimation of post-fertilization cell cycle and cell cycle duration

In theory, the number of cells doubles per cell division. Therefore, Log2 (number of total cells per embryo measured in Fig EV3A) was used as the number of post-fertilization cell cycles in medaka embryos (Fig EV3B). The estimated number of cell cycles was further used to estimate the cell cycle duration (Fig EV3C).

### Western blot

Dechorionated embryos were homogenized by gentile pipetting up and down in ice-cold PBS in microtubes. After centrifugation, the supernatant was removed, and the pellets were snap frozen in liquid nitrogen and stored in a freezer. Samples were boiled with Laemmli sample buffer at 95°C for 5 minutes, run on SDS-polyacrylamide gels, and transferred to PVDF membranes (Immobilon-FL, Millipore, IPFL00010). Membranes were incubated with blocking buffer (Intercept blocking buffer, LI-COR, 927-60001) for 1 hour at room temperature, incubated with primary antibody overnight at 4°C (1/2000 × antibodies, 0.1 % Tween-20 in blocking buffer), washed with TBST, incubated with secondary antibody (1/10000 × IRDye, 0.1% Tween-20, 0.01% SDS in blocking buffer) for 1 hour at room temperature, washed with TBST, and air dried for a few hours at room temperature in the dark. Imaging was performed with Odyssey CLx (LI-COR). Band identification and measurement of signal intensities were performed using Image Studio (LI-COR). Antibodies used for immunofluorescence are listed in Table EV2. Chameleon Duo Pre-stained Protein Ladder was used as the size marker (LI-COR, 928-60000).

For quantitative western blot, linearity of signal intensity was first confirmed using titrated samples as a standard (Fig EV2D, E, EV5E, F). H3K9me3 intensity was then normalized by GAPDH intensity (Fig 2E, F, EV5G).

### Statistical analysis

For two-sample statistical tests, normality and equal variances were first tested using the Shapiro-Wilk test and the F-test, respectively, with p-value = 0.05. If both null hypotheses were not rejected, a two-sided unpaired Student’s t-test was performed. If the null hypothesis of F-test alone was rejected, we performed two-sided Welch’s t-test. Otherwise, two-sided Wilcoxon rank-sum test was performed. *** p < 0.001, ** p < 0.01, * p < 0.05, NS: not significant, respectively.

## Data availability

This study includes no data deposited in external repositories.

## Acknowledgements

We acknowledge all the laboratory members for everyday discussion and for their continuous support. We acknowledge Ryohei Nakamura (University of Tokyo) for critical reading of the manuscript and valuable discussion. This work was supported by JSPS KAKENHI Grant No. JP23K14121 to H.S.F., by JSPS KAKENHI Grant No. JP22K20625 and Grant No. JP23K14190 to T.I., by the World-leading Innovative Graduate Program for Life Science and Technology, by the Ministry of Education, Culture, Sports, Science and Technology to S.I., and by Japan Agency for Medical Research and Development (AMED) under Grant No. JP18gm1110007h0001 to H.T.

## Author Contributions

**Hiroto S Fukushima**: Conceptualization; funding acquisition; investigation; methodology; writing – original draft; writing – review & editing. **Takafumi Ikeda**: Investigation; funding acquisition. **Shinra Ikeda**: Methodology. **Hiroyuki Takeda**: Funding acquisition; supervision; writing – review & editing.

## Conflicts of interest

The authors declare no competing interests.

